# Activity based checkpoints ensure circuit stability in the olfactory system

**DOI:** 10.1101/156372

**Authors:** Kiely N James, Benjamin T Throesch, Weston Davini, Kevin T Eade, Sulagna Ghosh, Sohyon Lee, Nina Torabi-Rander, Kristin K Baldwin

## Abstract

Olfactory circuits function at birth, yet are continuously remodeled through the integration of adult-born interneurons into the olfactory bulb in a manner that preserves olfactory perceptual stability throughout adult life. The mechanisms that ensure appropriate circuit stability in this dynamic context remain poorly understood. Since interneurons sculpt the excitatory output of mitral and tufted (MT) neurons to the olfactory cortex, we predicted that MT neurons might instruct interneuron wiring in the adult brain. By blocking synaptic transmission from MT neurons we show that MT neuronal activity is critical to maintain olfactory bulb integrity and interneuron survival. Blocking interneuron death uncovered a second activity-dependent checkpoint regulating dendrite branching. In contrast, cortical circuits and MT neurons remain stable in the face of these silent and degenerating olfactory circuits. These studies identify a circuit-specific role for non-sensory activity in regulating integration of neurons into the adult brain, as predicted by previous computational models.

## Introduction

Neuronal circuits must adapt to changing environments and new developmental stages while preserving sufficiently invariant patterns of connectivity to maintain stable percepts, memories and innate behavioral responses. In sensory systems such as vision, hearing and touch, intrinsic genetic mechanisms govern the initial formation of neural circuits, while neural activity and competition between neighboring neurons carrying similar sensory information act to refine circuit architecture. This typically occurs during restricted critical periods after which circuit structures stabilize ^1,2^. Compared to other sensory systems, less is known about how neural activity influences the formation, structure and stability of successive olfactory processing circuits.

Olfactory sensory information is detected and processed by ensembles of neurons that exhibit varying degrees of spatial stereotypy. In the nose, olfactory sensory neurons (OSNs) bearing one of ˜1000 odorant receptors are distributed widely throughout the epithelium without spatial order. Yet, axons of similar OSNs converge in the olfactory bulb (OB) to form spatially invariant synaptic structures, termed glomeruli, which exhibit activity that depends on the odorant receptor expressed by the related class of OSNs. This forms a spatial map of odor information in the OB. Each glomerulus is innervated by a set of ˜20-50 mitral and tufted neurons (MT neurons) that project axons to multiple cortical processing centers. In these centers, much of the glomerular spatial map is discarded, perhaps most completely in the piriform cortex where wiring is predicted to be stochastic ^3-5^. These higher representations of odor information are thought to be influenced by the local interneuron circuits in the OB, although the mechanisms shaping these local circuits are incompletely understood.

An unusual feature of the olfactory system is that the local circuits in the OB are highly dynamic throughout the lifetime of an animal. OSNs are repopulated continually and form synapses with MT neurons as well as with periglomerular inhibitory interneurons. MT neurons extend lateral dendrites where they form reciprocal synapses with dendrites of inhibitory neurons in the OB, the majority of which are the granule cells (GCs) that are thought to sharpen the OBs representation of odors. GCs and most other interneuron populations in the OB are continuously generated throughout adult life. These adult-born neurons arise in the subventricular zone (SVZ) and migrate through the rostral migratory stream (RMS) into the OB where they move radially, extend dendrites and establish new synaptic connections with MT neurons. As an apparent counterbalance to this proliferation, a percentage of existing and new adult-born OB interneurons undergo programmed cell death. This unusual form of cellular plasticity must somehow be regulated to allow for consistent odor recognition over the lifetime of an animal.

The integration of new GCs is impacted subtly by sensory activity. Reducing OSN activity or altering GC intrinsic responsiveness to excitatory inputs reduces GC survival and slightly decreases dendritic length, while increased activity can result in slightly increased survival ^6-10^. However, in these situations, the overall OB architecture is maintained, suggesting a limited role for sensory inputs in sculpting gross scale OB circuitry. In addition, MT neurons fire action potentials in the absence of OSN input, either due to spontaneous intrinsic activity or in response to cortico-bulbar inputs ^11^. Whether this sensory-independent MT neuron activity plays a role in shaping or maintaining OB circuitry has not been addressed.

Here, we compare the role of activity in OSNs with that of MT neurons in the formation and maintenance of neural circuits in the OB and olfactory cortical regions. To accomplish this, we developed parallel genetic strategies for cell type-specific expression of the tetanus toxin light chain (TeNT).Expressing TeNT in neurons leads to cleavage of Synaptobrevin2/VAMP2, severely reducing synaptic vesicular release. We show that while TeNT expression in OSNs has only a modest effect on OB circuitry, inhibiting vesicular release from MT neurons results in dramatic and progressive degeneration and disorganization in the OB that is largely accounted for by disrupted interneuron maturation and survival. Cell autonomous loss of NMDA receptors in subsets of GCs also perturbs GC maturation in a normal OB. Blocking cell death only partially rescues GC maturation, uncovering an activity-dependent checkpoint for GC maturation that could serve to recruit adult-born interneurons to MT neurons with increased levels of activity. Finally, we show that the gross scale architecture of olfactory cortex and MT neuronal survival is largely preserved in both conditions. While not previously shown experimentally, this activity-dependent selection mechanism confirms predictions by computational models aimed at understanding olfactory coding stability in the context of continued adult neurogenesis ^12^

## Results

### Genetic strategy to inhibit vesicular release selectively in OSNs or MT neurons

To assess the impact of neurotransmission by OSNs or MT neurons on olfactory processing circuits, we employed a genetic strategy to block vesicular release in each of these neuronal populations (Fig 1A). One way to impair neurotransmission is to cleave the VAMP protein by expressing the Tetanus toxin light chain (TeNT) in neurons. Here we employ a mouse line in which expression of Cre recombinase (Cre) induces the TeNT-GFP fusion protein, which cleaves VAMP and impairs synaptic vesicular release13. To restrict the TeNT protein to OSNs we crossed the TeNT mouse line to the *OMP*-TENT ^14^. In parallel,we crossed the TeNT line to the *Pcdh21*-IRES-Cre mouse line, in which Cre expression in the OB is restricted to MT neurons (MT-T mice). These mouse lines allow us to compare the roles that OSN and MT neuron excitatory input play in shaping OB circuitry.

**Figure 1.**
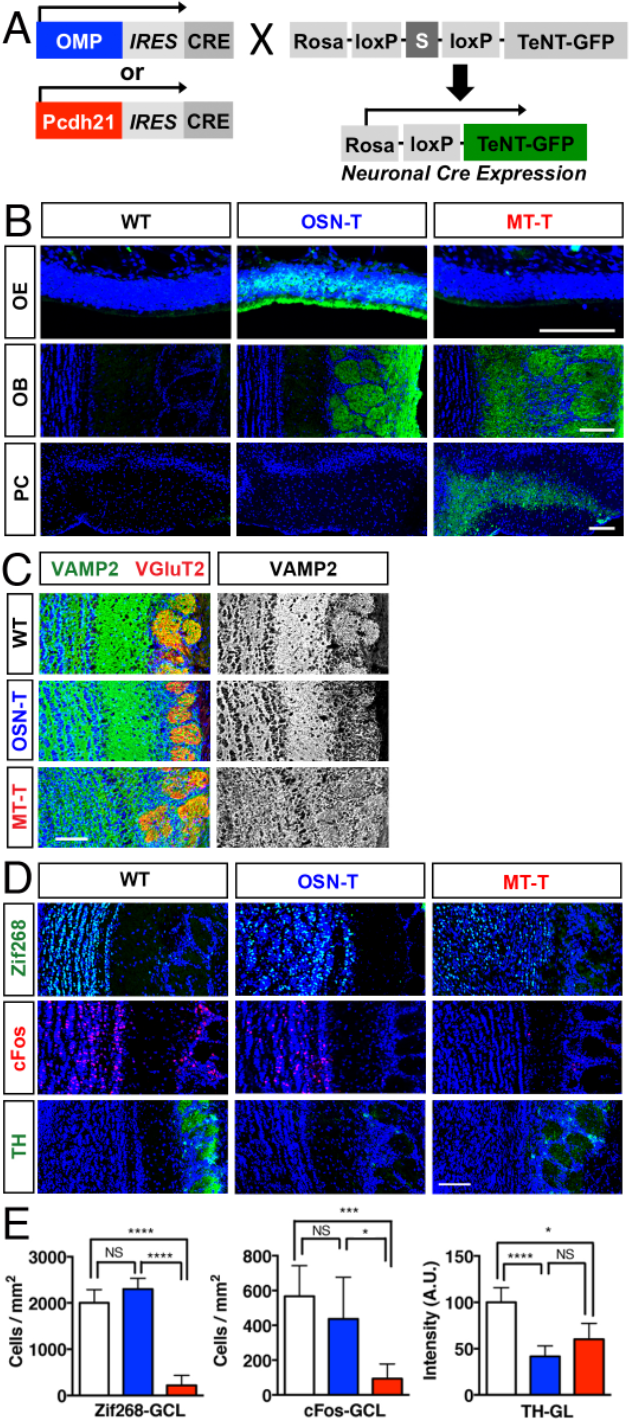
Genetic blockade of vesicular release in OSNs and MT cells and reduced neural activity in postsynaptic neurons. **A.** Genetic schema for generation of OSN-T or MT-T mice. **B**. TeNT-GFP in the OSN-T line is expressed in OSN cell bodies (OE) and axons (glomerular layer, OB) but not in PC. TeNT-GFP in the MT-T line is expressed in the MCL, EPL and GL, consistent with MT cell dendritic targeting, and in layer 2 of PC. Scale bars, 50 μm OE, 100 μm OB and CX. **C**. Decreased VAMP2 expression in the GL for OSN-T, and in the EPL and possibly GL for MT-T. **D.** Immediate-early genes Zif268 and cFos are reduced in the GCL of MT-T but not OSN-T, TH is reduced in the GL of both. Scale bar, 100 μm. **E.** Quantification of effects in D. Zif268: n= 6(WT), 3(OSN-T) and 5(MT-T) animals, 3-4 sections per animal. cFos: n= 4(WT), 2(OSN-T) and 3(MT-T) animals, 2-3 sections per animal. TH:n= 8(WT), 4(OSN-T) and 3(MT-T) animals, 2-4 sections per animal). Error bars indicate 95% confidence intervals. *p<.05, ***p<.001; ****p<.0001.

### Expression of TeNT is specific, cleaves VAMP and reduces immediate early gene expression in postsynaptic neurons

Cre expression in OSNs or MT neurons from OSN-T and MT-T mice should result in TeNT-GFP expression. Immunostaining showed that OSN-T mice expressed GFP in the OSN cell bodies and in their axonal terminals in the glomerular layer of the OB, but not elsewhere in the OB or brain (Fig 1B). Similarly, in MT-T mice GFP is present in the glomerular layer (GL) and external plexiform layer (EPL), where MT neuron dendrites project and form both pre- and postsynaptic terminals, in MT neuron cell bodies (MT layer of the OB), their axonal tract (LOT; lateral olfactory tract) and the layers where the MT axons terminate in olfactory cortex, but not elsewhere. Similar expression patterns are observed with the Ai9 Cre reporter line ^3,15^. These data confirm that for both crosses, the TeNT-GFP protein is present in the appropriate locations to cleave the VAMP protein and reduce synaptic transmission. We also observed that VAMP protein levels were reduced in the glomerular layer of OSN-T mice and in the EPL of MT-T mice, indicating that TeNT-dependent VAMP cleavage occurs in the correct locations to perturb neurotransmission (Fig 1C).

In the OB, sensory activity increases the expression of immediate early genes in postsynaptic target neurons. Blockade of OSN neurotransmission reduces but does not eliminate MT neuronal activity, so the effects of OSN vs. MT neurotransmission blockade are expected to differ ^11^. To examine the extent to which loss of VAMP in OSNs or MT neurons impairs activity levels in the granule cell neurons of the OB, we quantified the expression of two immediate early genes, Zif268/Egr1 and cFos. In the MT-T OB, we observed a significant decrease in both Zif268/Egr1 and cFos expression in the granule cells (Figs 1D, 1E). In OSN-T mice, however, Zif268 and cFos expression is indistinguishable from wild-type controls. These results suggest that for GCs, OSN neurotransmission blockade is less severe than MT neurotransmission blockade, perhaps due to the spontaneous and/or top down activation of MT neurons that persist absent OSN input ^11^.

We also examined a direct target of OSNs and MT neurons in the OB, the periglomerular neurons that express tyrosine hydroxylase (TH) in an activity dependent manner ^16^. We observed substantially reduced TH expression in the GL of OSN-T mice, as has been seen in other models of OSN signaling attenuation ^17-19,^ as well as in MT-T mice (Figs 1D, 1E).

### Blocking neurotransmission from MT neurons perturbs the OB circuitry differently than an OSN blockade

Previous studies have shown that a blockade of OSN input causes a modest reduction in the size of the OB but no change in its overall architecture ^18,20^. We also observe a reduction in the size of the OB in OSN-T mice (Figs 2B and S1A, S1B). In contrast, the OBs of adult MT-T mice are significantly smaller than those in the OSN-T mice and exhibit an aberrant laminar structure. Staining for MT neurons (PGP9.5) and OSN terminals (VGlut2) showed that the MT layer and the EPL were unevenly laminated and reduced in size, while the OSN glomerular layer was expanded (Fig 2A). These effects are not evident at birth but become detectable by postnatal day 11 (Figs 2B, 2C and S1C). As the animals age, both OSN-T and MT-T mice exhibit smaller OBs than in WT mice, but the OSN-T mice largely preserve the relative proportions of each layer while MT-T mice do not. At P125 MT-T mice exhibit an ˜80% reduction in the area of the granule cell layer, an irregular but substantial loss of the EPL and a concomitant enlargement of the MT and glomerular layers (Figs 2B, 2C). Strikingly, the MT-T OB shrank in absolute terms between P11 and P125. We found that the overall OB size difference could be largely attributed to differences in EPL and granule cell areas, while the area of the glomerular layer area was unchanged.

**Figure 2.**
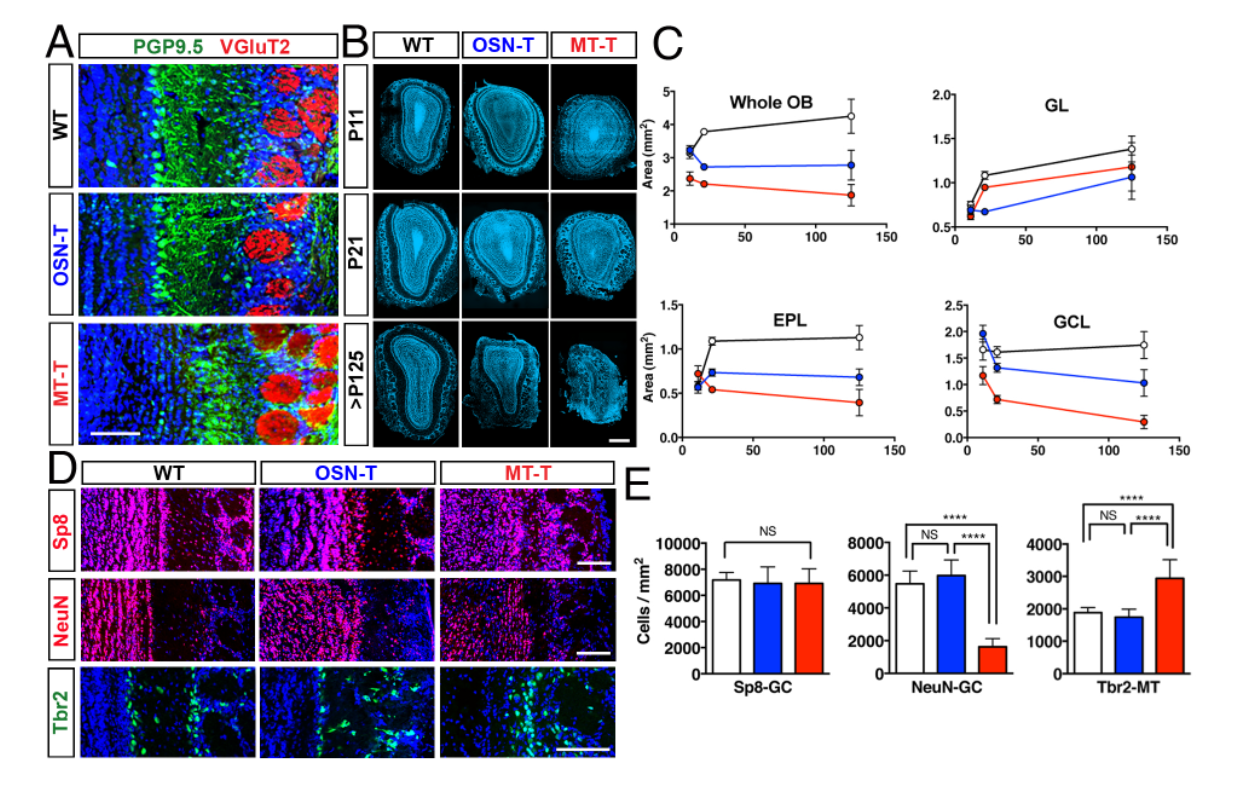
OB morphology and interneuron survival depend on MT cell vesicular release to a greater extent than OSN release. **A**. Laminar disorganization in P21 OB for MT-T but not OSN-T. PGP9.5 labels MT cells, VGlut2 labels OSN axons in the GL. Scale bar, 100 μm. **B, C**. Areal measurements of three OB layers reveal graded loss in OSN-T and MT-T, due to loss of GCL and EPL area. P11: n=3(WT), 2(OSN-T) and 3(MT-T); P21: n=4(WT), 6(OSN-T) and 4(MT-T); P125-137: n=2(WT), 1(OSN-T) and 2(MT-T) animals, 3 sections per animal. Scale bars, 500 μm. **D**. Cell density is unchanged in OSN-T, selectively decreased in MT-T for some neuronal populations. **E.** Quantification of D. Sp8 n=6(WT), 2(OSN-T) and 4(MT-T), 2-3 sections per animal. NeuN n= 5(WT), 3(OSN-T) and 4(MT-T) animals, 2-3 sections per animal. Tbr2-labeled MT cells are present in normal numbers in OSN-T and MT-T n=7(WT), 3(OSN-T) and 6(MT-T) animals, 2-4 sections per animal. Scale bars, 100 μm. Error bars indicate 95% confidence intervals. See also Figure S1.

For a more comprehensive picture of the aberrant circuit architecture in the OB of MT-T mice, we examined which cell types in the OB were lost in each mouse line (Figs 2D, 2E, S1D, S1E). The granule cells and most other inhibitory interneurons of the OB are continually generated throughout adult life and therefore consist of immature and mature populations. Two GC subtypes (5T4- and CR-expressing) were reduced in number in the MT-T but not OSN-T mice (Figs S1D, S1E). Other interneuron populations showed mixed responses to OSN-T and MT-T silencing, but in no case did we see greater interneuron survival in the MT-T compared to OSN-T line (Figs S1D, S1E). Both immature and mature GCs express Sp8, while only mature GCs express NeuN. In MT-T mice the density of Sp8 positive GCs was not changed, but the density of mature NeuN+ GCs was significantly reduced (Figs 2D, 2E), suggesting that MT activity blockade impairs GC maturation and/or survival.

### MT neurons persist in OBs with reduced sensory input and when they are prevented from communicating with postsynaptic targets

MT neurons and granule cells form reciprocal synapses, forming an intricate spatiotemporal web of excitation and inhibition. Given the sensitivity of inhibitory interneurons to reduction in their presynaptic inputs, we wished to assess the roles of vesicular release from OSNs and MT neurons in the maintenance of MT cells themselves. In the context of the pronounced morphological dysfunction that we observe in the MT-T OB, we hypothesized that the MT cells would undergo increased rates of cell death, either in order to maintain homeostatic signaling or due to loss of other factors that could promote survival. However, to our surprise, we saw no change in MT neurons (marked by Tbr2 ^21^) in OSN-T mice, and we observed an increase in Tbr2 cell density in MT-T animals (Figs 2F, and S1D), which most likely reflects normal survival of MT cells in a compressed EPL, since there is no evidence for production of “new” MT neurons in BrdU experiments that labeled postnatally generated neurons (Fig 3C).

**Figure 3.**
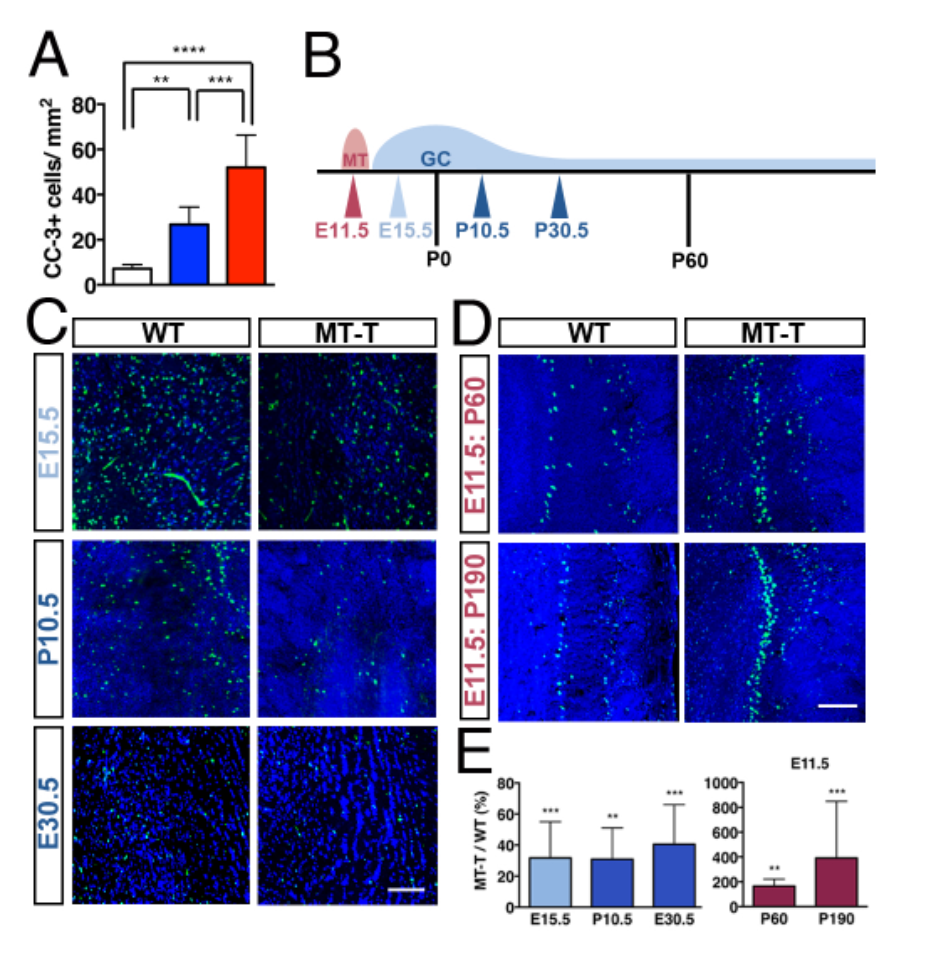
Long-term survival of embryonic and adult-born GCs requires MT cell vesicular release, but MT cells survive within an abnormal circuit. **A**. Graded increase in expression of cleaved caspase-3, a marker for cell death, in OSN-T and MT-T. n=9(WT), 5(OSN-T) and 5(MT-T), 3-5 sections per animal. Error bars indicate 95% confidence intervals. *p<.05; **p<.01; ****p<.0001. **B**. Schema of embryonic and postnatal BrdU injections. Red and blue curves indicate patterns of cell birth for MT cells and GCs respectively. Arrowheads indicate BrdU injection timepoints for MT cell (E11.5) and GC (E15.5, P10.5, P30.5) respectively. **C**. Embryonically and post-natally derived GCs are lost at equal rates in MT-T mice. Scale bar, 100 μm. **D**. Presumptive MT cells survive into advanced adulthood in MT-T mice. Scale bar, 100 μm. **E**. Quantification of C and D; MT-T GCs (blue bars) were reduced in density by 60-65% compared to WT for all three birthdates. BrdU staining in GCL at >P125 (E15.5 injection: n=2(WT), 5(MT-T) animals, 3-4 sections per animal; E10.5 injection: n=2(WT), 3(MT-T) animals, 2-6 sections per animal; E30.5 injection: n=2(WT), 3(MT-T) animals, 2-3 sections per animal). MT cells (maroon bars) are spared in MT-T OB. BrdU staining in MCL/EPL at P60 or P137 (P60: n=1(WT), 1(MT-T) animals, 4 sections per animal; P137:n=4(WT), 3(MT-T), 2-3 sections per animal. *p<.05; **p<.01; ***p<.001; ****p<.0001. See also Figure S2.

### Inhibitory neurons are produced and migrate properly but die in the MT-T OB

The loss of granule cells and other OB interneurons could be due to an effect on their precursors in the RMS. We performed several assays to examine precursor proliferation and early migration. Ki67, a marker of proliferating cells, was expressed at similar density in the RMS in each line (Figs S2A, S2B). Likewise, RMS immunoreactivity for Nestin, which labels neuronal precursors, was comparable in WT and MT-T mice. Finally, we counted BrdU-labeled cells in the RMS 24 hours after injection (Figs S2A, S2B). Using this method we again saw no difference in the number of newly generated cells between WT and MT-T mice. To directly examine the number of dying cells in the RMS we stained for cleaved caspase-3 (CC-3), a marker of apoptotic cells. The density of CC-3+ cells in the RMS of mice of each genotype was equivalent (Figs S2C, S2D). In contrast, the granule cell layer of the OB showed a nearly 10-fold increase in CC-3 cell density in MT-T mice, and a small but detectable increase in OSN-T mice (Fig 3A). These results show that the production and survival of OB interneuron precursors does not depend on MT neurotransmission and support a role for MT neurotransmission at later stages of postnatally-born GC maturation.

In addition to postnatally-born GCs, the olfactory bulb also contains a population of embryonically-born GCs. These neurons are of interest because they mature and begin to form synapses with MT neurons prior to birth ^22^ suggesting that they might be less sensitive to loss of activity than postnatally-born GCs ^6^ To address this possibility, we birthdated GCs by BrdU labeling at an embryonic timepoint (E15.5) or at two postnatal timepoints (P10.5 or P30.5) (Fig 3B). We quantified BrdU+ cell density in the GCL for all groups at P125-P135. Surprisingly, we found that embryonic and postnatally-born GCs were equally sensitive to loss of MT input, (E15.5; 68% reduction, P10.5; 69% reduction, P30.5; 59% reduction) (Figs E). In contrast, BrdU-birthdated MT neurons showed significant increases in MCL BrdU+ cell density (Figs 3D, 3E), while postnatal BrdU injections did not label MT cell layer neurons (Fig 3C). These data confirm previous reports that MT neurons are not replaced postnatally and establish that their survival does not depend on communication with their postsynaptic target neurons or on the preservation of normal OB circuit organization.

### Granule cell morphology is aberrant in MT-T mice

The development of postnatally-born GCs occurs in stages ^23^. Once GCs reach the OB, they first extend a single dendritic branch that reaches toward but does not enter the EPL (Stage 3). Next the dendrite enters the EPL and GCs begin to produce action potentials (Stage 4). Finally the dendrites become increasingly branched in the EPL and dendritic spines emerge (Stage 5). Previous studies demonstrated that under chronic naris occlusion, GCs exhibit a modest decrease in spine density and either a small decrease ^10^ or no change ^24^ in total dendritic length or number of branches. Given the relative magnitude of cell death and loss of GCL and EPL regions, we hypothesized that in MT-T mice individual GCs might exhibit more pronounced changes in morphology compared to GCs in sensory-silenced models. To compare the features of GC neurons in MT-T, OSN-T and WT mice, we engineered two lentiviral tracers and labeled postnatally-born GCs via intraventricular injection. First we labeled GCs with a lentivirus expressing GFP under the human *UbC* promoter (Figs S3A, S3C). Examination of GCs from P21 mice revealed a dramatic reduction in dendritic complexity in MT-T mice but not in OSN-T mice (Fig 4A).

**Figure 4.**
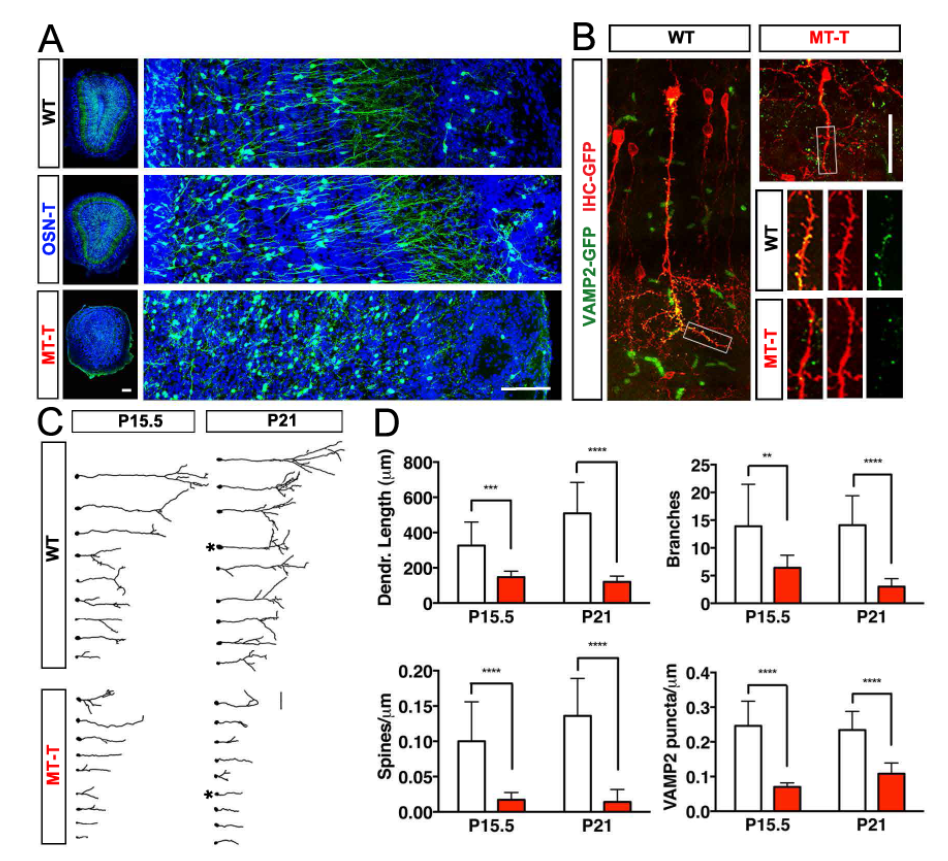
When MT cell vesicular release is inhibited, granule cell neurites are less complex and form fewer synapses. **A** Intraventricular injection of lenti-GFP reveals morphological deficits in MT-T GCs. Scale bars, 100 μm**.B** Representative range of GC reconstructions revealed by intraventricular injection of lenti-VAMP2-Venus fusion protein. Neurons within each group are arranged in descending order of total dendritic length. Starred neurons are shown in panel C. Scale bar, 50 µm. **C**. Examples of lenti-VAMP2-Venus labeled P21 GCs. Boxed dendritic branches are inset. Scale bar, 50 µ m. **D**. Quantification of morphological and synaptic features of reconstructed GCs. White bars indicate WT, red bars indicate MT-T. P15.5: n=14(WT), 37(MT-T); P20.5: n=9(WT), 15(MT-T) from 3 animals. Error bars indicate 95%confidence intervals. ***p<.001; ****p<.0001. See also Figure S3.

To more closely examine the structural aberrations in the GCs of MT-T mice, we also produced a lentiviral vector that encodes a fusion of VAMP2 and Venus, a variant of GFP (Fig S2B). In neurons expressing this virus, GFP is targeted to synaptic sites and accumulates in puncta that are easily quantified. To visualize the entire neuron, we also stained the cells with an antibody to GFP, which labels the entire dendritic tree. This staining was conducted in a different channel (red; Fig 4B). Three-dimensional reconstructions revealed that the GCs of MT-T mice exhibit drastically reduced total dendritic length and complexity (number of branches) as early as P15.5. Furthermore, both spine density and number of VAMP containing puncta were dramatically reduced (Figs 4B-4D). These studies show that the surviving GCs in MT-T mice exhibit a perturbed morphology resembling that of immature GCs in WT animals.

### Blocking cell death uncovers a developmental checkpoint regulated by MT activity

One hallmark of programmed cell death in neurons is neuritic degeneration, in which dendrites and axons shorten prior to cell death (reviewed in ^25^. We wondered if the small, simplified dendritic trees that we observed in activity-attenuated GCs were a symptom of imminent cell death or instead, a sign that these neurons had never matured. To distinguish between these models we used a Bax knockout (KO) to block apoptosis ^26^. The Bax genetic deletion was bred into the TeNT and *Pcdh21*-IRES-Cre mouse lines, which were then crossed to generate MT-T Bax KO mice. These mice exhibited dramatically reduced density of CC-3+ cells in the GCL, showing that loss of Bax rescues the GC cell death phenotype (Figs 5C, S4E). In addition, the size of the OB in MT-T Bax KO was partially rescued due to an increase in the GCL area and a modest increase in EPL area (Figs 5A, 5B).

**Figure 5.**
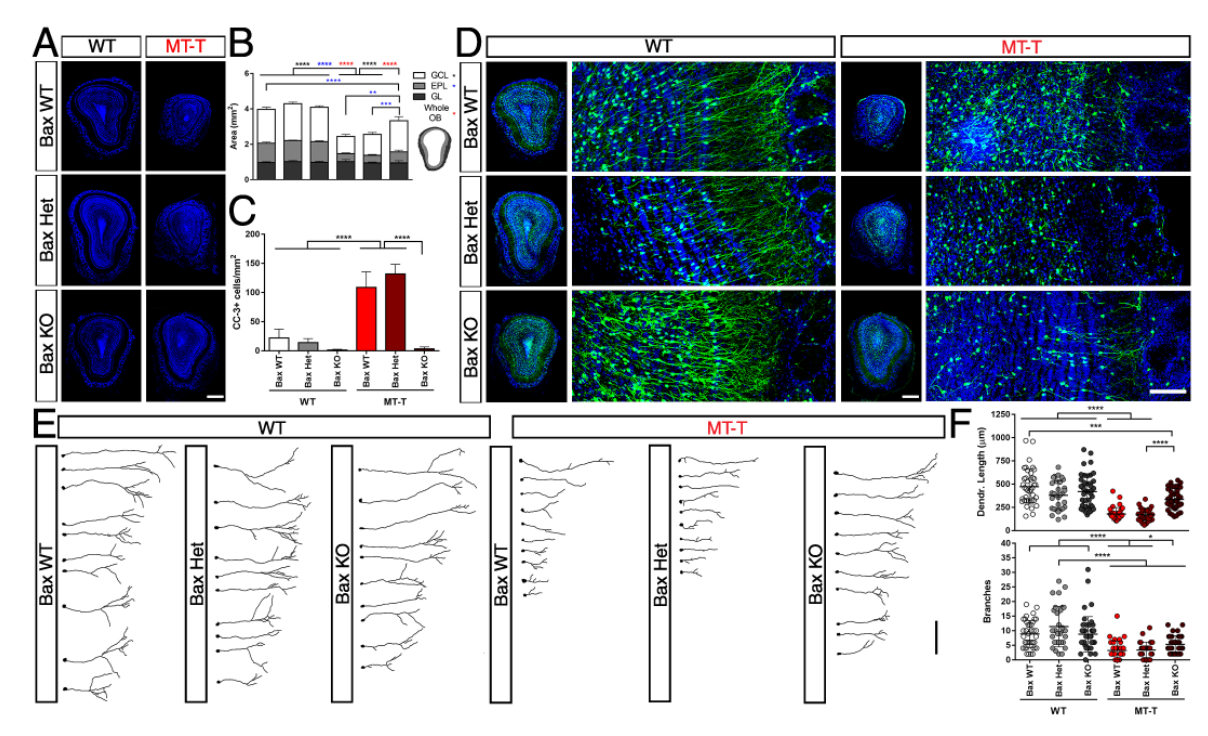
Inhibiting apoptosis in GCs with Bax KO mice partially rescues their morphological defects under MT cell vesicular blockade. **A, B.** Areal measurements of three layers reveals Bax KO partially rescues OB size in MT-T mice, due to increases in GCL and EPL areas. P21: n=5 animals per genotype, 3 slices per animal. Scale bar, 500 μm. Error bars indicate 95% confidence intervals. No difference in GL area. Red asterisks indicate significance for the whole OB area, blue for the EPL area, and black for the GCL area. **C.** Bax KO ameliorates expression of cleaved-caspase 3 in MT-T mice. n=5 animals per genotype, 3 slices per animal. **D.** Lentivirally labeled GCs expressing HA-Flag-GFP exhibit morphological deficits in MT-T GCs. Scale bars 500 μm, 100 μm. **E.** Representative reconstructions of lenti-HA-Flag-GFP labeled p21 GCs. Scale bar, 100 μm. **F.** Quantification of morphological features of reconstructed GCs. n=40 (WT, Bax WT), 31 (WT, Bax Het), 43 (WT, Bax KO), 34 (MT-T, Bax WT), 32 (MT-T, Bax Het), 40 (MT-T,Bax KO) from 4 animals. *p<0.05; **p<0.01; ***p<0.001; ****p<0.001.

Having validated our cell death blockade, we assessed its impact on GC maturation by sparsely labeling newborn GCs using a lentiviral vector encoding HA-tagged GFP (Fig 5D). Three-dimensional reconstructions show GCs in MT-T Bax KO mice are not fully mature (Fig 5E). Blocking apoptosis allows these neurons to extend dendrites further than in the MT-T mice, but their dendritic branching remains significantly reduced compared to GCs in WT mice, consistent with the partial rescue of EPL area that we observe (Figs 5E, 5F).

To confirm this result in a WT OB background, we used lentiviral vectors to block apoptosis in sparsely labeled GCs either by over-expressing Bcl2 or by expressing p53 shRNA. Each virus also contained GFP to mark the neurons (Fig S4A). We did not detect any rescue of the GC branching, dendrite length or spine number with either manipulation (Fig S4B-D). These results suggest that non-sensory activity levels gate both GC survival and morphological maturation.

### Granule cell morphological development depends on NMDA receptors and depolarization

The data we report are consistent with a model in which the reduction in excitatory input to GCs is greater in MT-T mice than in OSN-T mice. An alternative model would be that rather than neurotransmitters, some other vesicularly-released signaling molecule are selectively blocked in the MT-T mice, leading to the extensive circuit degeneration we observe. To differentiate between these possibilities we devised two strategies to reduce input to GCs without perturbing MT neurons

First, we established a genetic approach to eliminate NMDA receptor-mediated input onto sparse GCs in an otherwise normal OB. Both NMDA and AMPA receptors are expressed at high levels by GCs ^27^ and contribute functionally to GC excitation by MT cells, with NMDA receptors specifically generating the slow component of the GC EPSC ^28^. We obtained mice in which Cre expression leads to a functional knockout of NR1, an obligate subunit of NMDA receptors ^29,30^. We performed P0 intraventricular injections of a lentiviral construct separately encoding Cre and GFP (lenti-Cre-GFP), mixed with a control lentiviral construct encoding HA-tagged GFP (Fig 5A, Figs S5A, S5C). Importantly, NMDA currents are found in GCs only after they migrate into the OB and begin to move radially to their final position ^23^. This finding excludes the possibility that eliminating NMDA receptor -dependent excitation could affect GCs during their migration through the RMS.

Three-dimensional reconstructions of control lenti-HA-GFP GCs and NR1-KO GCs at P21 revealed substantial reductions in total dendritic length, number of branches, and dendritic spine density only in the NR1 KO GCs (Fig 6A-C). The impact of NR1 deletion on GCs resembled that of the MT blockade, but was not as severe. This intermediate effect could be due to the fact that AMPAR-mediated excitation of GCs is preserved in NR1-KO GCs, whereas in the GCs of MT-T mice, both types of excitation are presumably decreased at MT-GC synapses.

**Figure 6.**
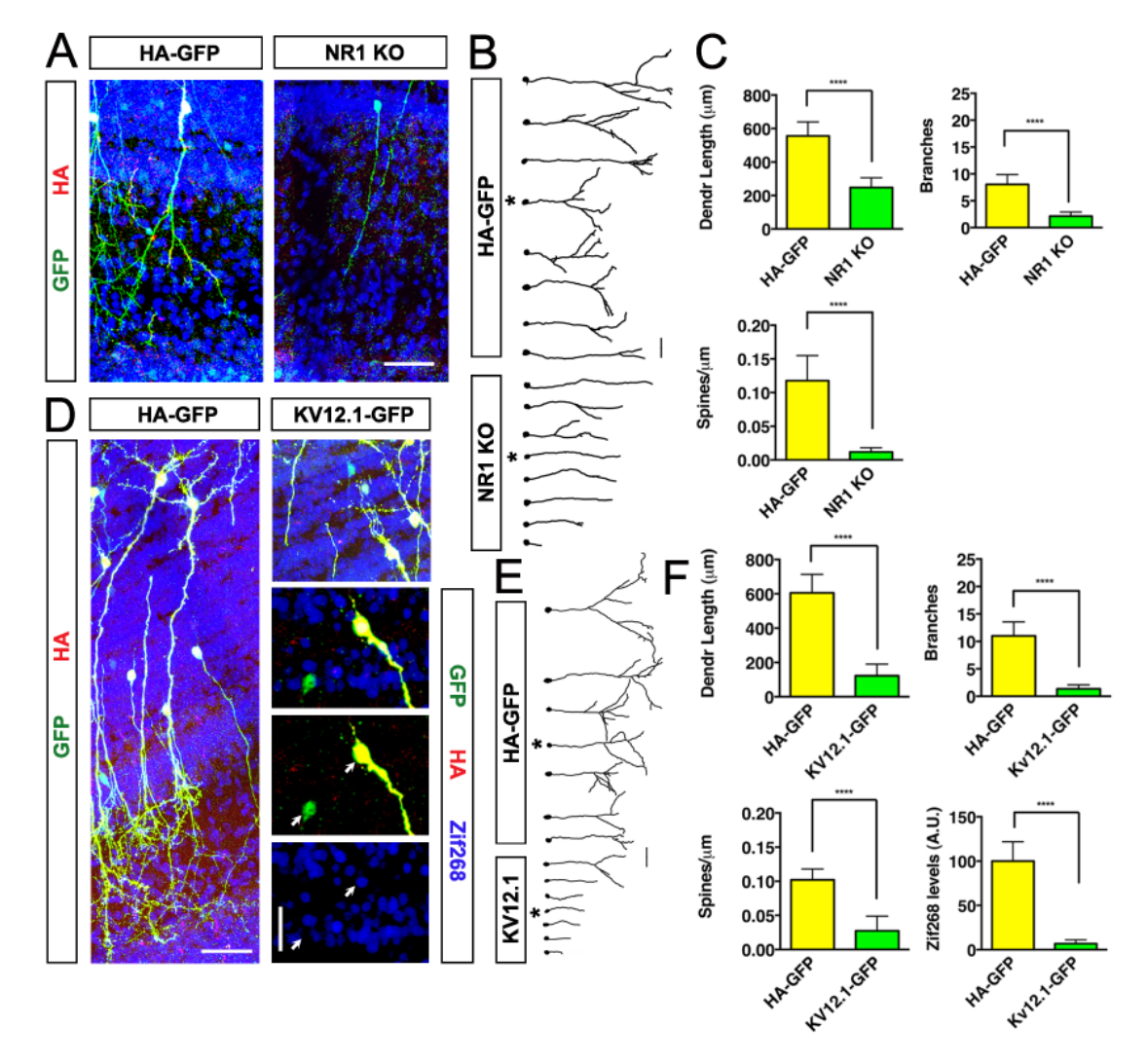
Two forms of activity blockade in sparse GCs recapitulate the morphological effects of loss of MT cell vesicular release. **A.** Examples of NR1 KO GCs or lenti-HA-GFP expressing GCs. Scale bar, 50 μm. **B.** Representative range of reconstructions of lenti-HA-GFP or NR1 KO GCs. Starred neurons are shown in panel D. Scale bar, 50 μm. **C.** Quantification of morphological features of reconstructed lenti-HA-GFP GCs or NR1 KO GCs. N=17(HA-GFP), 17(NR1 KO) from 3 animals. Error bars indicate 95% confidence intervals. ****p<.0001. **D.** Left and top right, examples of GCs expressing Kv12.1-SCPGFP or HA-GFP. Scale bar, 50 μm. Lower right, Zif268 levels in lenti-Kv12.1-SCPGFP GCs are drastically reduced compared to lenti-Zif268 levels in HA-GFP GCs. N=40 (lenti-HA-GFP), 10 (lenti-Kv12.1-SCPGFP) cells, 8 sections, 2 animals. Error bars indicate 95% confidence intervals. **E.** Representative range of GC reconstructions of lenti-HA-GFP or lenti-Kv12.1-SCPGFP GCs. Starred neurons are shown in panel A. Scale bar, 50 μm. **F.** Quantification of morphological features of reconstructed GCs expressing lenti-HA-GFP or lenti-Kv12.1-SCPGFP. N=11 (lenti-HA-GFP), 15 (lenti-Kv12.1-SCPGFP). Error bars indicate 95% confidence intervals. See also Figure S5.

We also sparsely infected GCs with a lentiviral construct expressing the voltage-gated potassium channel Kv12.1 (*Elk1, Kcnh8*), which begins to activate at -90 mV ^31^. Kv12.1 was cloned in frame with a fusion of the self-cleaving 2A peptide and GFP, allowing detection of Kv12.1-expressing neurons (lenti-Kv12.1-SCPGFP) (Fig S5B). Because the lentiviral construct drives gene expression using the synapsin promoter, expression levels in infected neurons should be very high compared to endogenous levels of Kv12.1 gene expression. As a result, infected neurons should be hyperpolarized relative to control neurons.

We injected a mixture of lenti-Kv12.1-SCPGFP and lenti-HA-GFP and examined the OB at P21 (Fig 6D). When we quantified levels of Zif268 expression in either group of infected neurons, we observed a dramatic decrease in Zif268 expression in Kv12.1-SCPGFP GCs relative to GFP-HA GCs (Figs 6D, 6F). The morphology of P21 Kv12.1-SCPGFP resembled that of GCs in the MT-T mouse and reconstructions revealed that all three measures of GC maturity were dramatically reduced Kv12.1-SCPGFP GCs relative to control GCs. (Figs 6E, 6F) This experiment provides additional evidence that the blockade of MT cell synaptic signaling in MT-T mice directly causes the death and morphological defects of GCs through a reduction in excitatory neurotransmission. Furthermore, it suggests that these effects are due to decreased GC activation (depolarization), as opposed to the loss of a synaptically localized signal such as cell-adhesion molecule mediated signaling.

### Blocking OSN or MT neurotransmission does not impact cortical circuit stability

The impact of sensory activity or MT vesicular release on cortical olfactory circuits has not been established. To address this issue, we first examined the AON and aPC to determine whether MT cell TeNT was functioning distally to reduce VAMP2 expression. VAMP2 reactivity was indeed reduced in layer 1 of AON and aPC, where MT axons synapse onto the dendrites of their postsynaptic partners (Fig 7A). Next, we examined the expression of Zif268 in each target and found, as expected, that levels were significantly reduced in the MT-T mice, with OSN-T mice exhibiting intermediate levels of reduction (Figs 7A, 7C). These data show that the MT-T and OSN blockade are detected at the level of the third order neurons in olfactory cortex.

**Figure 7.**
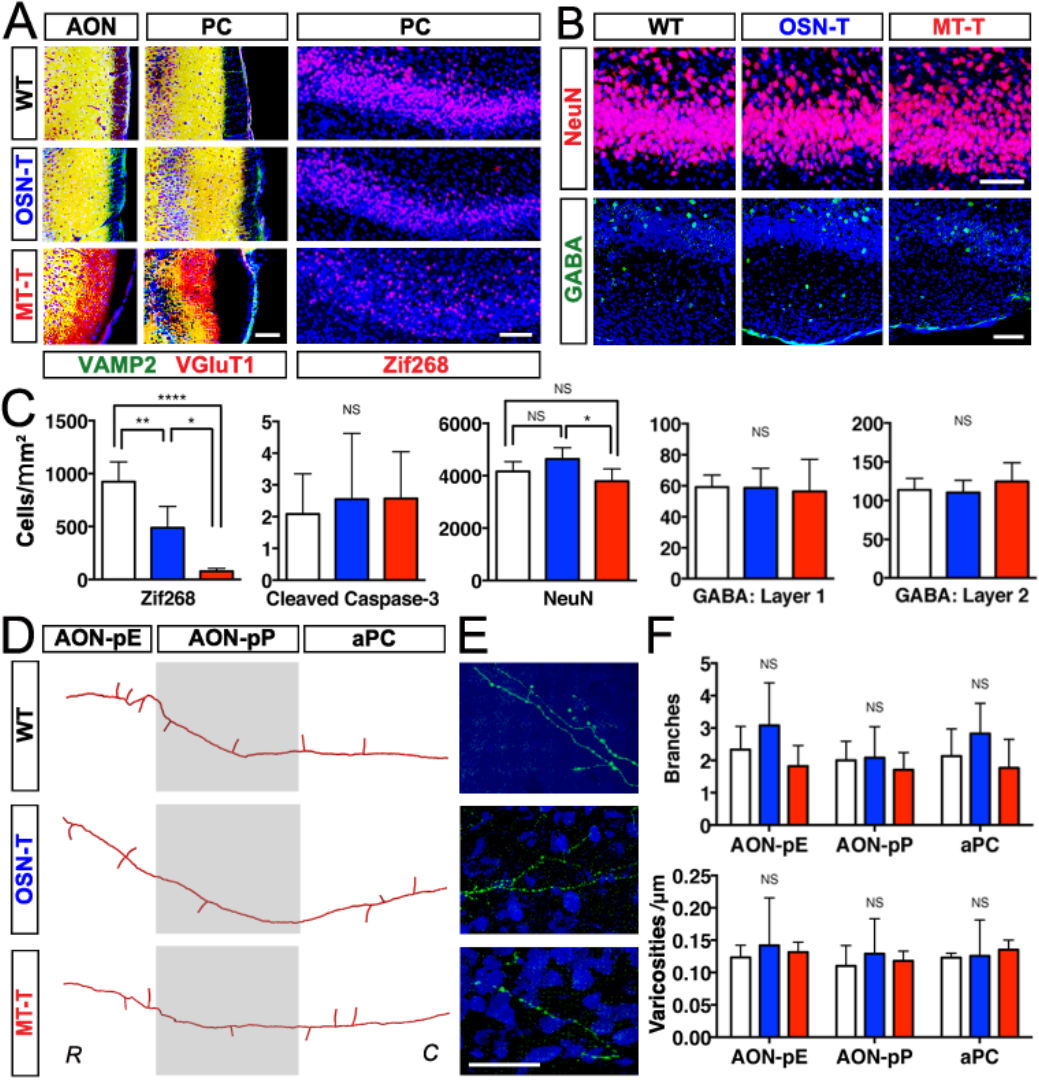
Axonal projections and target neurons in olfactory cortical regions are stable when sensory input or MT vesicular release is reduced. Left panel, VAMP2 levels are reduced in layer 1 of AON and aPC in MT-T mice. Scale bar, 100 μm. Right panel, Zif268 levels are intermediate in OSN-T PC and low in MT-T PC. Scale bar, 100 μm. **B**. Target neurons are present at normal densities in OSN-T and MT-T PC. Scale bar, 100 μm. **C**. Left, quantification of Zif268 levels. n= 6(WT), 3(OSN-T) and 5(MT-T) animals, 3-4 sections per animal. Next, target neurons in piriform cortex do not die at increased rates, as assessed with cleaved caspase-3 staining. N=8(WT), 2(OSN-T) and 5(MT-T) animals, 1-8 sections per animal. Number of cells counted= 62(WT), 22(OSN-T) and 21(MT-T). Final three panels, quantification of NeuN and GABA data shown in B. NeuN: n=6(WT), 3(OSN-T) and 3(MT-T) animals, 4-9 sections per animal. GABA: n= 8(WT), 4(OSN-T), and 5(MT-T), 3-5 sections per animal. Error bars indicate 95% confidence intervals. **D**. Representative reconstructions of MT cell axonal projections in three target areas of cortex. R: rostral, C: caudal. **E.** Images of higher order MT cell axonal branches in piriform cortex. Note varicosities. Scale bar, 50 μm. **F.** Primary branching and varicosity density are unaffected in OSN-T and MT-T mice. Quantification of number of primary branches: n= 10(WT), 8(OSN-T) and 12(MT-T) axons analyzed from n=(WT), 4(OSN-T) and 5(MT-T) animals. Quantification of varicosity density: n= 4-5 animals per genotype, 3 sections/area/animal. Error bars indicate 95% confidence intervals. *p<.05, **p<.01; ****p<.0001. See also Figure S6.

Next, we asked whether reduced neurotransmission from MT neurons would lead to apoptosis in target neurons in the cortex, as it does in the OB. Staining for neuronal cell death using CC-3 revealed no detectable increase in MT-T or OSN-T mice (Fig 7C). Similarly, the numbers of NeuN and GABA positive inhibitory interneurons in layers 1 and 2 of the piriform cortex were not significantly different from wild type either mouse line (Figs 7B, 7C). Taken together, these results provide evidence for stability of postsynaptic targets of MT neurons both when sensory input is blocked (OSN-T) or when all direct excitatory input from MT neurons is blocked (MT-T).

In the auditory system, reducing presynaptic sensory input to projection neurons can restrict their axonal branching patterns ^32^. To assess the role of neurotransmission in regulating the primary branching patterns of MT cell axons, we labeled sparse MT neurons using Sindbis virus that we engineered to express fluorescent proteins ^3,^ (Fig S5A). Three dimensional reconstructions of MT axonal projections into AON and aPC in both OSN-T and MT-T mice revealed no detectable changes in primary branch location, primary branch number or layer ramification (Figs 7D, 7F). We also used sparse Sindbis labeling to examine MT dendrite structure in the OB. In OSN-T mice, MT cells were qualitatively indistinguishable from those of WT littermates, with obliquely angled lateral branches, a single, thick apical dendritic branch whose tuft evenly filled a single glomerulus as in WT OBs (Fig S5B). The MT cells of MT-T mice had somewhat aberrant morphologies, but they maintained an apical tuft that extended into a single glomerulus. These data suggest that MT neurons depend neither on OSN input nor on interaction with postsynaptic neurons to correctly innervate a single glomerulus.

Reduction of sensory inputs to MT neurons or blockade of vesicular release in MT neurons might be expected to alter their formation of presynaptic structures. We tested this hypothesis by counting the morphologically distinct axonal varicosities found on neurons labeled with Sindbis, which we have previously shown to be localized with presynaptic markers ^3,33^. MT cell presynaptic varicosity density is similar for all three cortical target regions in WT mice (Figs 7E, 7F). Thus, in contrast to the drastic effects of blocking OSN- or MT-cell vesicular release within the OB, both the cells of the cortical target regions and the gross branching and presynaptic structures of MT axons appear resistant to these manipulations.

### Discussion

Here we delineate the differential impact of the two major sources of excitatory neurotransmission on odor processing circuits of the OB and piriform cortex. While the gross scale architecture of MT cells and their target regions in the piriform cortex and AON are preserved when either MT cell or OSN neurotransmission to these regions is reduced, we have uncovered an unexpectedly large impact of MT cell activity on the stability of OB circuits. The loss of circuit architecture under MT cell silencing derives from increased loss of previously integrated neurons and, more acutely, from a severe impairment of later-born interneuron survival and dendritic development. These results extend our understanding of olfactory circuit development, maintenance and function and provide insights into mechanisms regulating the integration and survival of neurons in adult circuits relevant for regenerative medicine.

### MT cells are resistant to circuit alteration or loss of activity

MT cells are essential for the sense of smell, comprising the only neuronal population that connects the nose to the cortex. Here we show that MT cell activity is essential for OB circuit assembly and maintenance. In contrast, MT cells themselves survive in normal numbers and project to cortical target areas, under sensory blockade (OSN-T), when their own activity is blocked and when many of their synaptic partners in the OB die (MT-T). The activity-*independent* survival and maturation of MT cells may be linked to the fact that they appear early in development—prior to the birth of most INs—and are subjected to widely varying levels and patterns of activity as the OB develops around them ^34,35^. These findings define a new instructive role for MT cells in building and maintaining olfactory circuit architecture. The stability of MT cells throughout various perturbations of circuit activity may help to preserve olfactory “resolution” or perceptual stability, particularly in light of the apparent stochastic patterns of wiring in the piriform cortex.

### An activity-based checkpoint for GC maturation and survival

MT cells exhibit a basal firing rate that is also affected by breathing or sniffing ^36^. Sensory activity, by contrast, is patterned and sparse, but thought to increase the level of MT firing above baseline ^37-39^. Here we show that the basal firing rate of MT cells absent sensory input is sufficient to permit the assembly and maintenance of a largely normal OB, consistent with other studies ^6,40,41^. In addition, using BrdU labeling we show that MT activity also governs the fate of previously integrated embryonically-born interneurons. To our knowledge, this is the first evidence that the survival of embryonically-born GCs can be influenced by afferent excitation.

These results suggest a model in which the survival and maturation of OB interneurons is regulated by an activity-based checkpoint. Checkpoints are used to control the rate and identity of incoming entities. Setting the threshold for proceeding through the checkpoint at the level of input produced by intrinsic activity in MT cells, could help to preserve an even overall pattern of GC wiring independent of a particular sensory environment. Indeed a published mathematical model of the dynamic OB circuitry showed that implementing an activity dependent survival alongside random cell death over time could result in maximal odor discrimination that is robust to a changing environment (Cecchi et al., 2001). Our study is the first to experimentally demonstrate that such a signal is provided globally to embryonically born and adult born interneurons through the basal levels of MT firing that are independent of the odor environment rather than through sparse or intermittent patterns of sensory neuron input.

### Dissecting the mechanisms of activity dependent GC maturation

A topic of interest in developmental neuroscience and regenerative medicine is to define the mechanisms and transcriptional programs that enable neurons to integrate into circuits, either early in development, later in adult life or in situations of neurodegeneration or injury. Here, we show that activity regulates two distinct steps of GC integration into a preexisting circuit. First, we show that GCs depend on excitatory neurotransmission for survival at an immature stage, both in a globally silent OB and when individual GCs in an otherwise normal OB lack NMDA receptors or are hyperpolarized through aberrant potassium channel expression. This suggests that one direct consequence of neurotransmission through NMDA receptors is to block transcriptional programs that induce apoptosis. Alternatively, this protective effect could be mediated post-translationally.

Next, by blocking cell death using the Bax knockout mouse (as well as by overexpressing Bcl2 and reducing p53 via shRNA) we show that rescue from cell death does not fully rescue maturation of GCs. Rather, the GCs extend longer primary dendrites and appear to regain some of their radial architecture but they do not extend lateral dendritic trees or achieve complete innervation of the EPL, despite the presence of MT cells in normal numbers. This shows that excitatory neurotransmission is required to enact or stabilize a transcriptional (or post-translational) program that regulates morphological and synaptic maturation. Interestingly, the observation that activity acts to promote the expansion of GC dendritic territory runs counter to observations of activity-dependent pruning or reduction of neurite territory in other sensory systems ^42-45^. This suggests that pathways regulating dendritic expansion vs. retraction/pruning may be differentially regulated by activity depending on the functional requirements of a particular circuit.

This study may provide insight into strategies aimed at replacing or maintain neurons in degenerating or injured brain regions. For example, we show that blocking apoptotic pathways can improve neuronal survival and lead to increased neuronal maturation in a partially silent adult circuit. This result suggests that blocking apoptosis until circuit activity is restored could be a useful therapeutic strategy, if other adult circuits are regulated similarly to the olfactory bulb. Future studies of this system will be required to identify the specific molecular pathways that lead to cell survival and dendritic maturation in the maturing and adult olfactory system.

## Author Contributions

KNJ and KKB conceived of experiments; KNJ, BT, WD, KTE, SG, SL and NTR carried out experiments; SG contributed reagents; KNJ, BT, WD, KTE, and SL analyzed data; KNJ and KKB wrote the paper.

## Acknowledgements

We thank Simon Pieraut for assistance with intraventricular injections, Kathy Spencer for imaging and analysis advice, and Anton Maximov for critical feedback. This work was supported by funding from the National Institute on Deafness and other Communication Disorders (DC012592 to K.K.B.), the National Institute of Mental Health (MH102698 to K.K.B.), the California Institute for Regenerative Medicine (RB3-02186 to K.K.B.), the Baxter Family, Norris and Del Webb Foundations (K.K.B.), by Las Patronas and the Dorris Neuroscience Center (K.K.B.).

## Experimental Procedures

### Mouse strains

TeNT-GFP mice were generously provided by Martin Goulding ^13^Zhang et al., 2008)^15^ *OMP*(^14^ and the TeNT*OMP*TeNT-IRES-CRE (OSN CRE) line was developed in the Axel laboratory (TeNTEggan et al., 200426-GFP positive; we observed no morphological or other differences between CRE and -GFP controls. TonegawaBax^29,30^Knudson et al., 1995)Tsien et al., 1996a Tsien et al., 1996b;; Jax stock number 005246. stock number 002994. Conditional NR1 knockout mice were generated in the laboratory of Susumu (Tsien et al., 1996a; Tsien et al., 1996b); Jax stock number 005246.

### Tissue preparation and immunohistochemistry

Mice P15 and younger were euthanized and brain or olfactory epithelial tissue were post-fixed for one hour (for cryosections) or overnight (for vibratomed sections) in 4% PFA at 4° C overnight. Mice P15 and older were transcardially perfused with 4% PFA in PBS containing CaCl2 and MgCl2, and brains were placed in 30% sucrose in PBS (for cryosections) or 4% PFA (for vibratomed sections) at 4° C overnight. For cryosections, brains were embedded in OCT, stored at -80° C and later sectioned at 25 µm with a Leica CM3050 S cryostat. Slides were allowed to air dry for ˜1 h, post-fixed for 10 minutes in 4% PFA, rinsed several times with PBS, and treated with blocking buffer (5% heat-inactivated horse serum in PBS-T (PBS + 0.1% TritonX-100)) for 1 h at room temperature. Primary antibodies, diluted at 1:500 (unless otherwise noted) in blocking buffer, were applied to slides and left on overnight at 4° C. Following three 10-minute rinses in PBS-T, secondary antibodies, diluted at 1:1000 in blocking buffer were applied for 1-2 h at room temperature. Slides were rinsed several times in PBS-T and mounted using ProLong Gold mounting media either with or without DAPI (Life Technologies P36935 or P36934). For vibratomed sections, brains were sectioned at 80 µm using a Leica VT1000S and stained, using the same reagents and timing as above, in 48 well plates. Primary antibodies used:

5T4 (rabbit, Abcam ab129058)

BrdU (1:400, mouse, BD 555627)

Calbindin (rabbit, Swant CB38)

Calretinin (goat, Millipore AB1550)

cFos (rabbit, Calbiochem PC38)

Cleaved caspase-3 (rabbit, CST 9661)

GABA (rabbit, Sigma-Aldrich A2052)

GFP (rabbit, Invitrogen A11122 or sheep, Serotec 4745-1051)

HA (mouse, Covance MMS-101P, BioLegend 901501)

Ki67 (1:300, rabbit, Acris DRM004)

PGP9.5 (rabbit, Abcam ab10404)

Nestin (mouse, MAB5326)

NeuN (mouse, Millipore MAB377)

Parvalbumin (goad, Swant PVG-214)

Sp8 (goat, SCBT C-18)

Tbr2 (rabbit, Millipore AB2283)

TH (rabbit, Pel-Freez P40101-0)

VAMP2/Synaptobrevin (mouse (69.1) Synaptic Systems 104211)

VGluT2 (GP, Millipore AB2251)

Zif268/Egr1 (rabbit, SCBT sc-189),

For BrdU staining, air-dried cryosections or vibratomed sections were rinsed briefly with water and 2N HCl, then incubated in 2N HCl for an hour at 37° C. Sections were then rinsed with PBS-T and processed normally beginning with the blocking step. For GFP staining of TeNT-GFP, an HRP-conjugated secondary antibody followed anti-GFP primary antibody application. After extensive washes, fluorescein-conjugated tyramide from a TSA kit (PerkinElmer NEL701A001KT) was applied to slides for ten minutes. Finally, after 2-3 additional hours of rinses, Sudan Black dye (0.1% in 70% EtOH) was applied for 10 m to reduce neuronal autofluorescence.

### BrdU administration

Bromodeoxyuridine was diluted to 3.125 or 12.5 ug/ul in PBS and injected intraperitoneally into individual mice or pregnant dams at 50 µg/g body weight. For the experiment examining recently born cells in the RMS, mice were injected once, 24 h prior to euthanization. For the granule cell survival experiments, mice or pregnant dams were injected twice, 24 h apart, and euthanized at later timepoints.

### Lentiviral constructs

Lentiviral constructs were cloned using modified versions of the FUGW shuttle vector ^46^Lois et al., 2002). Unless otherwise noted, they encode proteins driven by the UbC promoter. Three constructs were gifts from A. Maximov: Lenti-eGFP, lenti-Kv12.1-SCP-GFP and lenti-VAMP2-Venus. Murine ^31^ gene expression is driven by a (Zhang et al., 2009^47^synapsinlentiviral promoterAddgene. In lenti-VAMP2-Venus, rat VAMP2 (whose amino acid sequence is 100% identical to murine VAMP2) and the GFP variant Venus are cloned in frame as a fusion protein, with Venus at the C-terminal. FLAG-HA-GFP was generated in laboratory of W. Harper (lentiviralSowa et al., 2009Addgene) and cloned into the lentiviral shuttle vector by K.N.J.; Addgene plasmid 22612. Lenti-CRE-GFP was generated in the laboratory of T. Jacks; Addgene plasmid 20781. Lenti-CRE-GFP encodes GFP driven by the UbC promoter and CRE driven by the PGK promoter.

### Production of lentivirus

Shuttle vectors were co-transfected with packaging vectors pCMVΔ8.9 and pVSVg, or pMD2.G, pRSV-Rev, and pRRE (generated in the laboratory of Didier Trono; Addgene plasmids 12259, 12253, and 12251) into HEK293T cells using the calcium phosphate method. Virus was collected at 48 h post-transfection, concentrated by ultracentrifugation (2h at 113,000 *g,* 4°C), resuspended in a small volume of PBS and stored at -80 °C.

### Intraventricular injection of lentivirus

A simple manual air pressure-based system using a 3 mL syringe, plastic adaptors and thin tubing and was constructed and fitted with a glass micropipette (˜10 µm tip diameter). Neonatal mice (P0 or P1) were anesthetized for several minutes on wet ice. The injection site was targeted by drawing a line from bregma (visible beneath the skin) to the top of each eye with a fine-tip marker, then crossing it perpendicularly approximately ⅓ of the way down from the top (bregma). At an injection depth of 1-2 µ m relative to the skin surface, a volume of 0.5-2 µl of virus was ejected. Mice were allowed to recover beneath a warm lamp for several minutes and then returned to the home cage.

### Olfactory bulb-targeted injection of Sindbis virus tracers

Injections were performed as previously described ^3^Ghosh et al., 2011) but without reference to a fluorescently labeled glomerulus. Briefly, 3-week-old mice were anesthetized with isoflurane (2% in 100% O2) and olfactory bulbs were injected with Sindbis virus encoding fluorescent proteins (GFP alone, RFP alone or GFP and RFP in a single construct). Mice were euthanized 48 h after injection.

### Image collection and image processing

Other than some MT cell/axon reconstructions, all images were acquired on a Nikon C2 or Nikon A1 confocal microscope, often using automated XY stitching. Image analysis was performed using Nikon NIS-Elements, MBF Neurolucida Explorer and Adobe Photoshop. Cell counts were either automated or done manually depending on the signal strength and background, and normalized to anatomically defined regions using DAPI counterstaining. Several MT cell/axon reconstructions were acquired on an Olympus FluoView 500 confocal microscope.

To calculate the area of each OB layer, we used maximal coronal sections through the olfactory bulb, just anterior to the AOB. Three concentric regions were drawn using DAPI staining as a guide. The first region followed the MCL and the area within it was assigned to the GCL. The second region followed the bottom edge of the GL. From that region’s area, the GCL area was subtracted and the resulting area was assigned to the EPL. Finally, a third region followed the top edge of the GL, excluding the ONL. From that region’s area, the EPL layer was subtracted and the resulting area was assigned to the GL.

For interneuron counts, the regions sampled were as follows. NeuN-GC and Sp8-GC: a superficial section of the GCL (excluding the MCL) extending to approximately [inline1] the radius of the OB and corresponding to an area ˜0.04 mm^2^. 5T4-GC and CR-GC: a superficial section of the GCL whose dimensions were chosen to best capture the staining pattern observed in WT animals, corresponding to an area of ˜0.055 mm^2^ (5T4) or ˜0.07 mm^2^ (CR). CR-PG and CB-PG: a section of the GL corresponding to an area of ˜0.1 mm^2^ (CR) or ˜0.065 mm^2^ (CB). Blanes(CB)-GCL: either the entire GCL or a section of it that was radially proportional to the whole. PV-EPL: a section of the EPL corresponding to an area of ˜0.1 mm^2^. Tbr2-MT: a section encompassing the MCL, EPL and GL corresponding to an area of ˜0.2 mm^2^. OB Cleaved caspase-3 staining was measured within the entire GCL.

For BrdU staining, experiments B2-B4 used the entire GCL, or a representative portion of it to exclude regions with artifactually high background. Experiment B1 used a section including the MCL, EPL and the basal portion of the GL that included BrdU+ cells.

For cortical reconstructions of mitral cell axons, images were acquired either on a Nikon C2 or an Olympus Fluoview 500 at 20x. Cells to trace were chosen using these criteria: 1) the cell body was large, triangular in shape and lay within the MCL 2) a thick apical dendrite ended in a tuft filling a single glomerulus 3) the axon projected posteriorly through the LOT and remained visible through anterior PC. Images acquired on the Olympus Fluoview 500 were manually stitched in XY space. For all data sets, successive z stacks were manually aligned to make a rostral-caudal superstack in Neurolucida. Brain regions (AONpe, AONpp, aPC) were identified using DAPI nuclear counterstaining. For primary branch analysis, the main axon was traced and branch points where secondary branches emerged were marked and traced for a short distance. For varicosity analysis, three representative images (each corresponding to a z-stack of a single section) per target brain region were analyzed for each animal. A maximal projection of each image was used to trace a stretch of secondary- or higher order MT cell axonal branch and count the number of corresponding varicosities.

For cortical cell density counts, the regions sampled were as follows. Zif268 and NeuN: layer 2 of PC. GABA: layer 1 and 2 of PC, separately.

### Neuronal reconstructions

Images of granule cells were acquired on either a Nikon C2 or Nikon A1 as automatically stitched large image z-stacks at 40x. For each image, up to three neurons were chosen to reconstruct, in increasing order of the distance of the soma from the mitral cell layer. Criteria for reconstruction were that the neuron appeared to be completely contained within the section/z stack with no obviously truncated branches at the top or bottom, and that the soma contain some detectable native GFP, indicating a minimum level of VAMP2-GFP expression. Using Neurolucida 6.0 (MBF Bioscience), we reconstructed the granule cell’s dendritic tuft, excluding any trailing dendrite. Dendritic length was measured for the entire dendritic tree, and the number of branches was calculated as one less the sum of dendritic nodes and terminals. Spines and (for VAMP2-Venus tracings) presumptive VAMP2 puncta (native GFP puncta on dendritic branches or spines) were marked and attached to the dendritic tracing for density analysis. Dendritic spines were marked separately, based on both red (immunostained GFP) and green (unstained GFP) channel signal.

### Statistical analysis

GraphPad Prism 6 and 7 were used to perform unpaired t-tests, One-way ANOVA, Two-way ANOVA and Tukey’s multiple comparisons tests.

All procedures were performed in accordance with the guidelines and standards of The Scripps Research Institute’s Animal Care and Use Committee.

